# Physical and mental health status in *Toxoplasma*-infected women before and three years after they learn about their infection: The role of Rh factor

**DOI:** 10.1101/141580

**Authors:** Blanka Šebánková, Jaroslav Flegr

**Author notes:** Corresponding author. Address: Division of Biology, Faculty of Science, Charles University in Prague, Vinicna 7, 128 44, Prague, Czech Republic. *E-mail address*: 1 (J. Flegr).

## Abstract

Latent toxoplasmosis is known to be associated with specific changes in animal and human behavior and human personality. Many toxoplasmosis-associated shifts, such as an extroversion-introversion shift or a trust -suspicion shift, go in opposite directions in men and women. The stress coping hypothesis suggests that such behavioral effects of toxoplasmosis are side effects of chronic stress caused by lifelong parasitosis and associated health disorders. Several studies have searched for, and typically found, indices of impaired health in infected subjects. However, subjects were always aware of their toxoplasmosis status, which could influence obtained data and cause false-positive results of the studies. Here we searched for differences in physical and mental health status among 39 *Toxoplasma*-infected and 40 *Toxoplasma*-free female university students who completed identical questionnaires (N-70, and anamnestic questionnaire), before and 3 years after they were informed of their toxoplasmosis status. Our results showed that infected women showed indices of poorer health status, not only after, but also before they were informed of their infection. In accordance with previously published data, these indices were more numerous and stronger in Rh-negative than in Rh-positive women. Present results suggest that observed indices of poorer health and symptoms of chronic stress in *Toxoplasma*-infected subjects are real. Due to its high (30%) prevalence, toxoplasmosis could represent an important factor for public health.

## 1. Introduction

The prevalence of toxoplasmosis in the human population varies between 5–80% in various countries, depending on temperature, humidity, hygienic standards and kitchen habits (Tenter et al., 2000; Pappas et al., 2009). It is decreasing in many European countries and in Northern America, however, it is mostly increasing in the most populous countries in Eastern Asia, like China. *Toxoplasma gondii* reproduces sexually in the intestinal cells of its definitive host (any feline species, including domestic cats) and asexually in bodies of intermediate hosts (all warm blooded animals, including humans). In infected immunocompetent humans, a short phase of acute toxoplasmosis, which is accompanied by many nonspecific clinical symptoms including high body temperature, headache, swollen lymphatic nodes and fatigue, spontaneously proceeds into a lifelong latent phase. Latent toxoplasmosis is characterized by the presence of anamnestic IgG anti-*Toxoplasma* antibodies in serum and slowly dividing bradyzoites in tissue cysts localized mainly in immunoprivileged organs such as the brain, eye, and testes. Latent toxoplasmosis is usually considered harmless for infected immunocompetent subjects. However, many observations – typically anecdotal – suggest that it increases the risk of various disorders, for recent review see (Flegr et al., 2014). Psychiatrists performed systematic studies of the clinical effects of toxoplasmosis. They showed that toxoplasmosis likely strongly increases the risk of schizophrenia, bipolar disorder, obsessive-compulsive disorder, as well as epilepsy and migraines. For reviews see (Flegr, 2015; Sutterland et al., 2015).

Latent toxoplasmosis is also known to be accompanied by many specific changes in behavior and personality of infected animal and human hosts. *Toxoplasma* is transmitted from an intermediate to a definitive host by predation. Therefore, the behavioral changes observed in infected animals, like that of humans, are usually considered to be a feature of the adaptive manipulation of the parasite, intended to increase the probability of transmission of the dormant stages of *Toxoplasma* from intermediate to definitive host by trophic route. However, based on the nature of observed changes, especially based on the fact that many *Toxoplasma*-associated changes are in opposite directions in men and women (Flegr, 2010), it has been suggested that they are just nonspecific side-effects of mild but continuous stress induced by the parasitic disease (Lindová et al., 2010). It is known that men and women behaviorally cope with chronic stress in opposite ways. Stressed women express increased extroversion; they are more likely to seek and provide social support (Stone and Neale, 1984; Rosario et al., 1988; Carver et al., 1989), join with others (Hobfoll et al., 1994), and verbalize towards others or the self (Tamres et al., 2002). In contrast, stressed men express decreased extroversion, and seem to use more individualistic and antisocial (e.g., aggressive, hostile) forms of coping (Carver et al., 1989; Hobfoll et al., 1994).

Several studies have tried to test the stress coping hypothesis by searching for indices of chronic stress in subjects with the latent form of *Toxoplasma* infection. For example, an ecological study performed in 2014 showed that the prevalence of toxoplasmosis in particular countries largely explains the variability in mortality and morbidity associated with many diseases and disorders (Flegr et al., 2014) between countries. Recently, a large cross-sectional study performed on a nonclinical cohort of 1,500 internet users showed that *Toxoplasma*-infected subjects scored more poorly in 28 of 29 monitored health-related variables than *Toxoplasma*-free subjects and reported significantly higher incidence of 77 of a list of 134 disorders reported by at least 10 participants of the study (Flegr and Escudero, 2016). The main problem of the last study was that its participants self-reported their toxoplasmosis status and therefore knew whether or not they were infected. Therefore, it can be speculated that at least some of *Toxoplasma-infected* participants reported more health problems because of autosuggestion, as they believed that *Toxoplasma* could (or must) negatively influence their health.

The aim of the present study, was to search for possible indices of poorer physical and mental health status in a group of past and present female university students. These students voluntarily participated in behavioral studies at the Faculty of Science, Charles University, and approximately three years later, also participated in economic game experiments at the University of Economics, Prague. Most participants filled in two identical questionnaires (an anamnestic questionnaire monitoring physical and mental health status, and a standard N-70 questionnaire monitoring seven potentially pathognomonic factors: anxiety, depression, obsession, hysteria, hypochondria, psychosomatic symptoms of vegetative lability, and psychasthenia) before and about three years after they were told whether they are *Toxoplasma*-infected or not. This experimental setup enables one to discriminate whether the *Toxoplasma*-infected subjects express the symptoms of impaired health because of the infection or because they know that they are infected and therefore believe that they should express (or report) such symptoms.

## 2. Material and Methods

### 2.1. Participants

Subjects were recruited from a pool of former biology students of the Faculty of Science, Charles University, Prague, who have voluntarily participated in various ethological studies over the past 20 years. All subjects were women who were tested for toxoplasmosis and rhesus (Rh) status between 2002–2013. These volunteers often take part in many studies, most of which are not related to *Toxoplasma* infection. Crucially, toxoplasmosis was not mentioned in the study recruitment invitation, and the experiments were run at the University of Economics—not the usual location for toxoplasmosis-related experiments. Moreover, the experimental games were primarily focused on the examination of the economic behavior of women (Lanchava et al., 2015). Therefore, it is unlikely that subjects suspected that the study was related to toxoplasmosis research. To achieve a balanced design and the highest possible power of the study, we invited all Rh-negative *Toxoplasma*-infected women in the pool (the rarest combination of the trait within the study), and also the same number of *Toxoplasma*-infected and *Toxoplasma*-free women. Therefore, the frequency of infected subjects and Rh-negative subjects did not correspond to their normal representation in the Czech population. All participants provided written informed consent. Subjects’ recruitment and data handling were performed in compliance with the Czech legislation in force, and were approved by the Institutional Review Boards of the Faculty of Science, Charles University (No. 2014/21).

### 2.2. Questionnaires

At the beginning of the present study, the volunteers were asked to complete an anamnestic questionnaire as well as a N-70 questionnaire. Most participants completed these questionnaires at two different time points: before testing for toxoplasmosis, and then about 3 years later at the beginning of the present study. In the anamnestic questionnaire, the participants rated their physical and mental health status by answering a panel of questions such as: “How often do you have allergies (or: non-allergic skin problems; digestive problems; common infectious diseases, such as flu; cardiovascular problems; orthopedic problems; neurological problems; headaches; other chronic or recurring pain; other chronic or recurring health problems; depression; other mental health problems; metabolic problems like diabetes; are you tired after returning from work/school; do you visit medical doctors (not dentists and not for prevention))?” using a 6-point Likert scale (0: never, 1: maximally once a year, 2: maximally once a month, 3: maximally once a week, 4: several times a week, 5: daily or several times daily). They were also asked how many times they used antibiotics in the past 3 years (5 means 5 or more times), and how many times they spent more than a week in the hospital in the past 5 years. The new version of the questionnaire completed after 3 years also contained the question “At what age do you expect to die?”.

The N-70 is originally a Czech questionnaire constructed for the assessment of seven areas of psychiatric symptom clusters - anxiety, depression, obsession, hysteria, hypochondria, psychosomatic symptoms of vegetative lability, and psychasthenia (Vacíř, 1973). The English version of the questionnaire is available at (Flegr et al., 2012). Subjects are asked to answer 70 questions using a 3-point agreement scale. Scores in each cluster range from 0–30. The total N-70 score is the sum for all 70 questions.

### 2.3. Serological tests

All testing for toxoplasmosis was performed at the National Reference Laboratory for Toxoplasmosis, National Institute of Public Health, Prague. The complement-fixation test (CFT), which determines the overall levels of IgM and IgG antibodies of particular specificity, and Enzyme-Linked Immunosorbent Assays (ELISA) (IgG ELISA: SEVAC, Prague) were used to detect *T. gondii* infection status of the subjects. ELISA assay cut-point values were established using positive and negative standards according to the manufacturer’s instructions. In CFT, the titre of antibodies against *T. gondii* in sera was measured in dilutions between 1:4 and 1:1024. The subjects with CFT titres between 1:8 and 1:128 were considered *T. gondii* infected. Only subjects with clearly positive or negative results of CFT and IgG ELISA tests were diagnosed as *T. gondii*-infected or *T. gondii*-free, whilst subjects with different results on these tests, or ambiguous results, were retested or excluded from the study.

A standard agglutination method was used for Rh factor examination. A constant amount of anti-D serum (human monoclonal antiD reagent; SeracloneH, Immucor Gamma Inc.) was added to a drop of blood on a white glass plate. Red cells of Rh-positive subjects were agglutinated within 2–5 minutes.

### 2.4. Data analysis

Statistica v. 10 was used for the statistical analysis. Differences in age were tested by a t-test. N-70 scores were analysed using an ANCOVA, with age as a confounding variable. Certain scores (obsession and, to a smaller extent, depression) had slightly asymmetric distributions; therefore, we repeated the analyses with nonparametric methods. However, results of the parametric and nonparametric analyses were qualitatively the same. Ordinal and binary data were analyzed by a partial Kendall’s correlation test, which is used to measure the strength and significance of the association among binary, ordinal, or continuous data regardless of their distributions and allows for the control of one confounding variable – in this case, age (Siegel and Castellan, 1988). The Excel spreadsheet used to compute the partial Kendall tau and the significance for variables A (diseases) and B (*Toxoplasma* infection), once C (age) was controlled based on Kendall Taus AB, AC, and BC-is available at: http://web.natur.cuni.cz/flegr/programy.php. The false discovery rate (preset to 0.1) was controlled with the Benjamini-Hochberg procedure (Benjamini and Hochberg, 1995). In contrast to the simple Bonferroni’s correction, this procedure also takes into account the distribution of p values of performed multiple tests.

All raw data are available as the Supporting Information S1.

## 3. Results

The final set contained 14 Rh-negative and 26 Rh-positive *Toxoplasma*-free women and 13 Rh-negative and 26 Rh-positive *Toxoplasma*-infected women. Infected women were older than *Toxoplasma*-free women (24.6 vs. 23.0, t_77_=−1.90, p=0.06). No significant effects of age (p=0.402, μ^2^=0.01), toxoplasmosis (p=0.206, μ^2^=0.02), and Rh factor (p=0.419, μ^2^=0.01) were observed, but a significant effect of toxoplasmosis-Rh factor interaction (p=0.011, μ^2^=0.08) on total N-70 score was detected with an ANCOVA. The same analysis was performed separately on the seven N-70 traits, and showed significant effects of toxoplasmosis-Rh factor interaction on obsession (p=0.036, μ_2_=0.06), vegetative lability (p=0.032, μ^2^=0.06), and psychasthenia (p=0.008, μ^2^=0.09). The visual inspection of Figure 1 suggests that the *Toxoplasma*-infected Rh-negative women had much higher N-70 traits, whilst *Toxoplama*-infected Rh positive subjects had approximately the same or slightly lower N-70 traits than corresponding *Toxoplasma*-free controls.

**Fig. 1.**
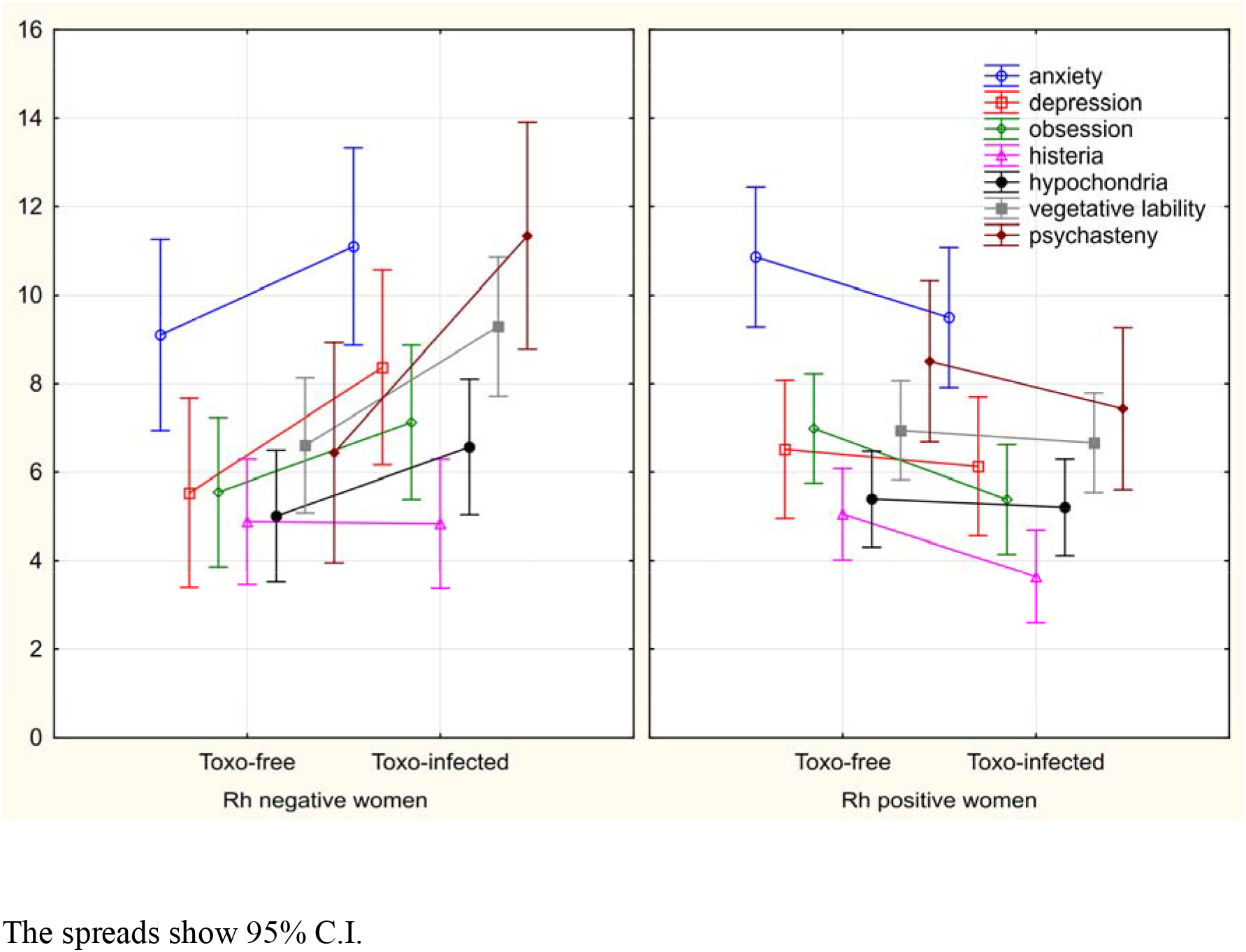
Differences between *Toxoplasma*-infected and *Toxoplasma*-free women in seven N-70 factors

The detailed examination of raw N-70 data showed that *Toxoplasma*-free women answered many questions of the N-70 questionnaire differently than *Toxoplasma*-infected women. Partial Kendall analyses, with age as a covariate, were performed on all 79 women, and showed that 13 of 70 observed differences remained significant after the correction for multiple tests. Separate analyses of 27 Rh-negative women and 52 Rh-positive women showed that the number of significant differences between *Toxoplasma*-infected women and *Toxoplasma*-free women was 32 of 70 and 12 of 70 in Rh-negative and Rh-positive women, respectively. While in Rh-negative women only 3 of 32 significant associations were negative, i.e., indicated better health status or wellbeing of *Toxoplasma*-infected women, in Rh-positive subjects, 11 of 12 associations, significant after the correction for multiple tests, were negative (see Tab. 1).

**Table 1.**
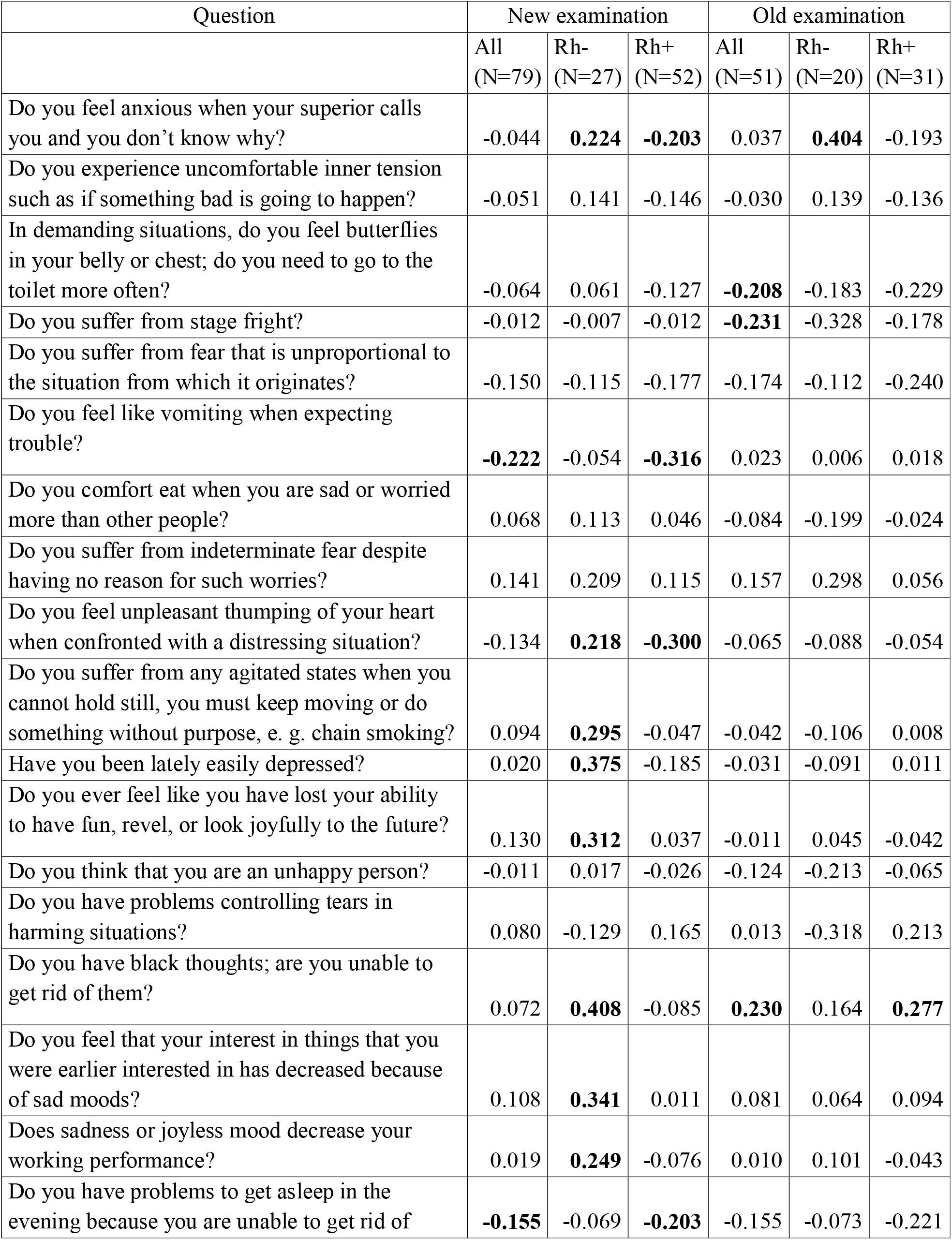
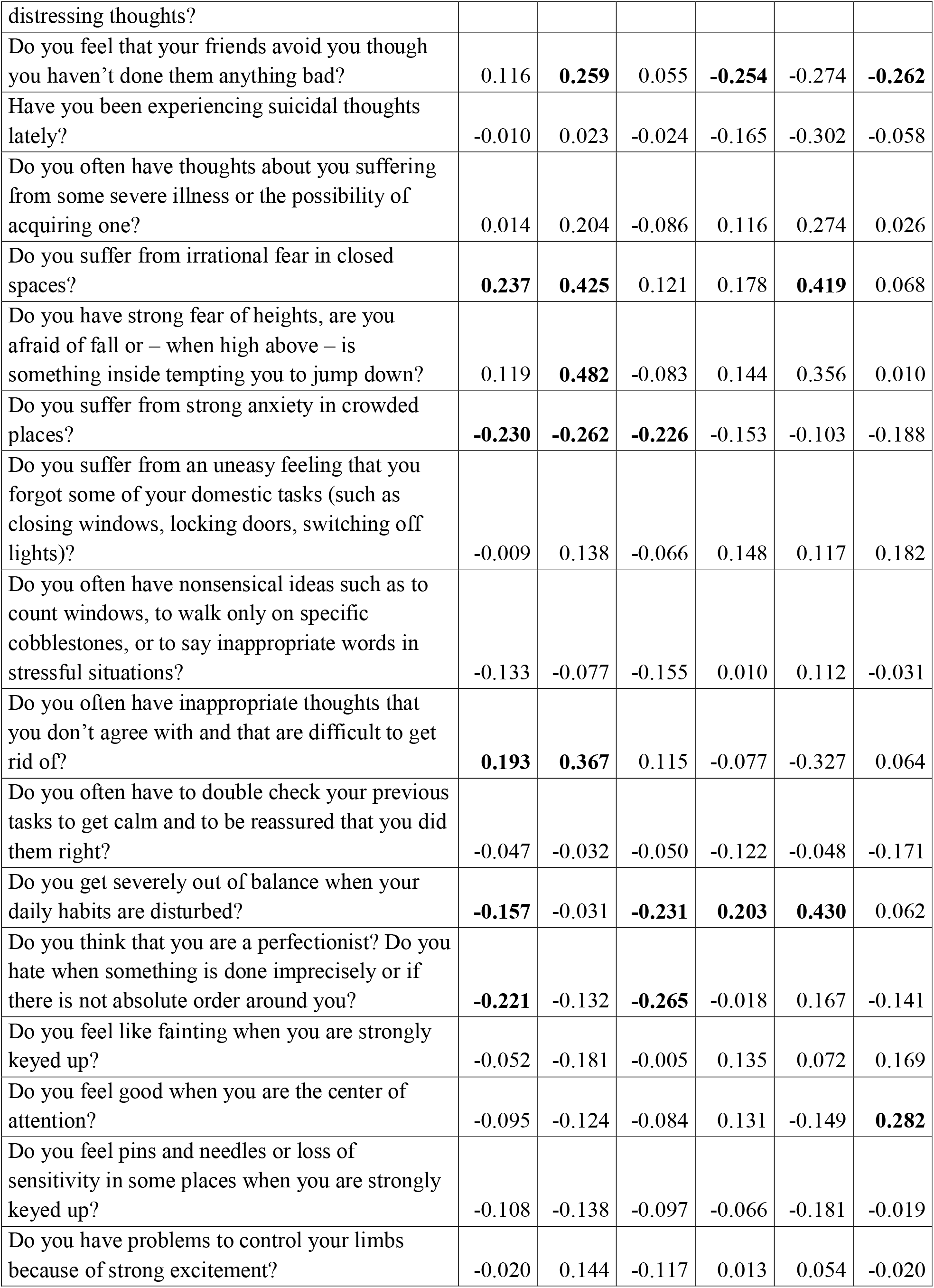
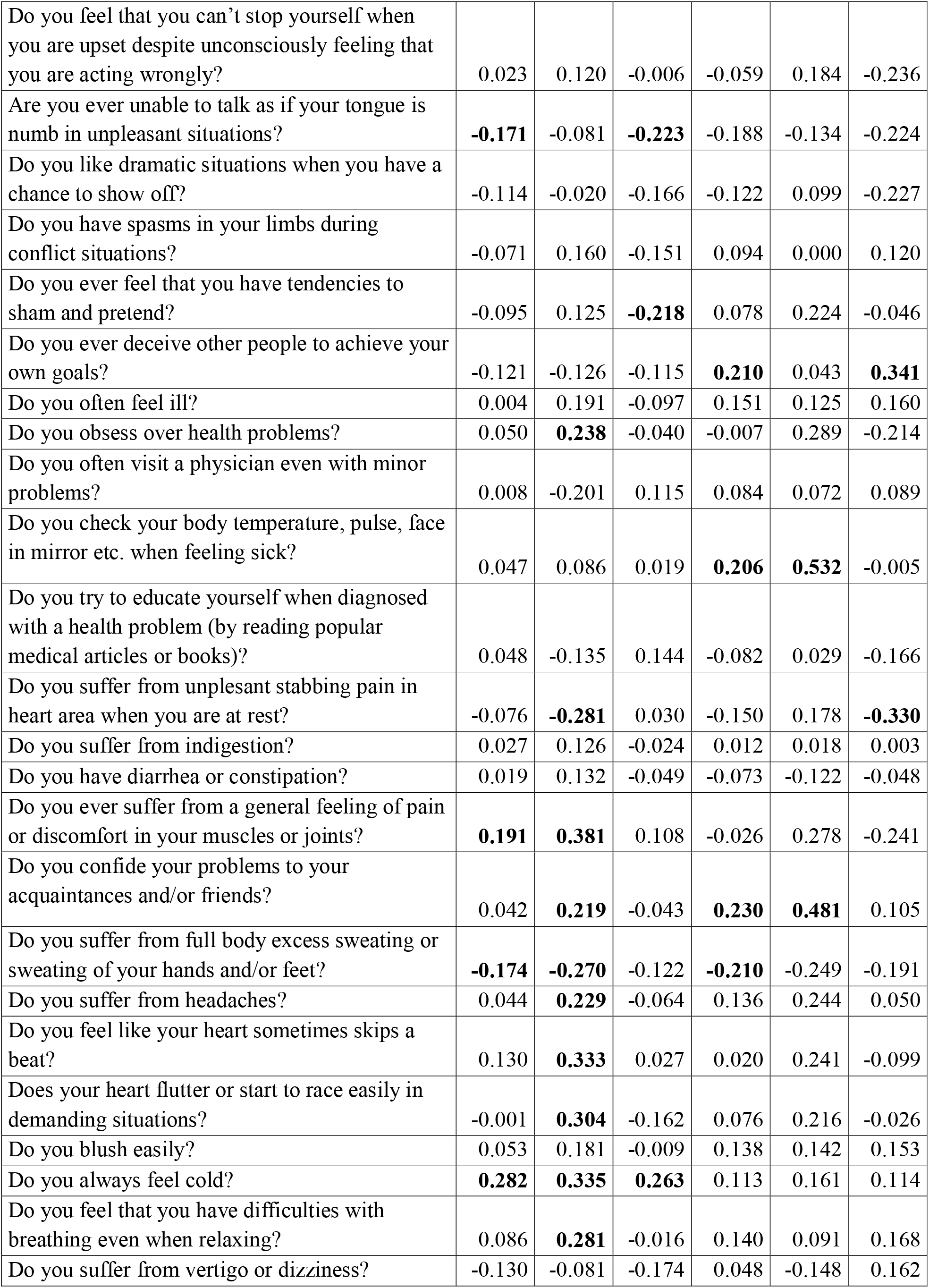
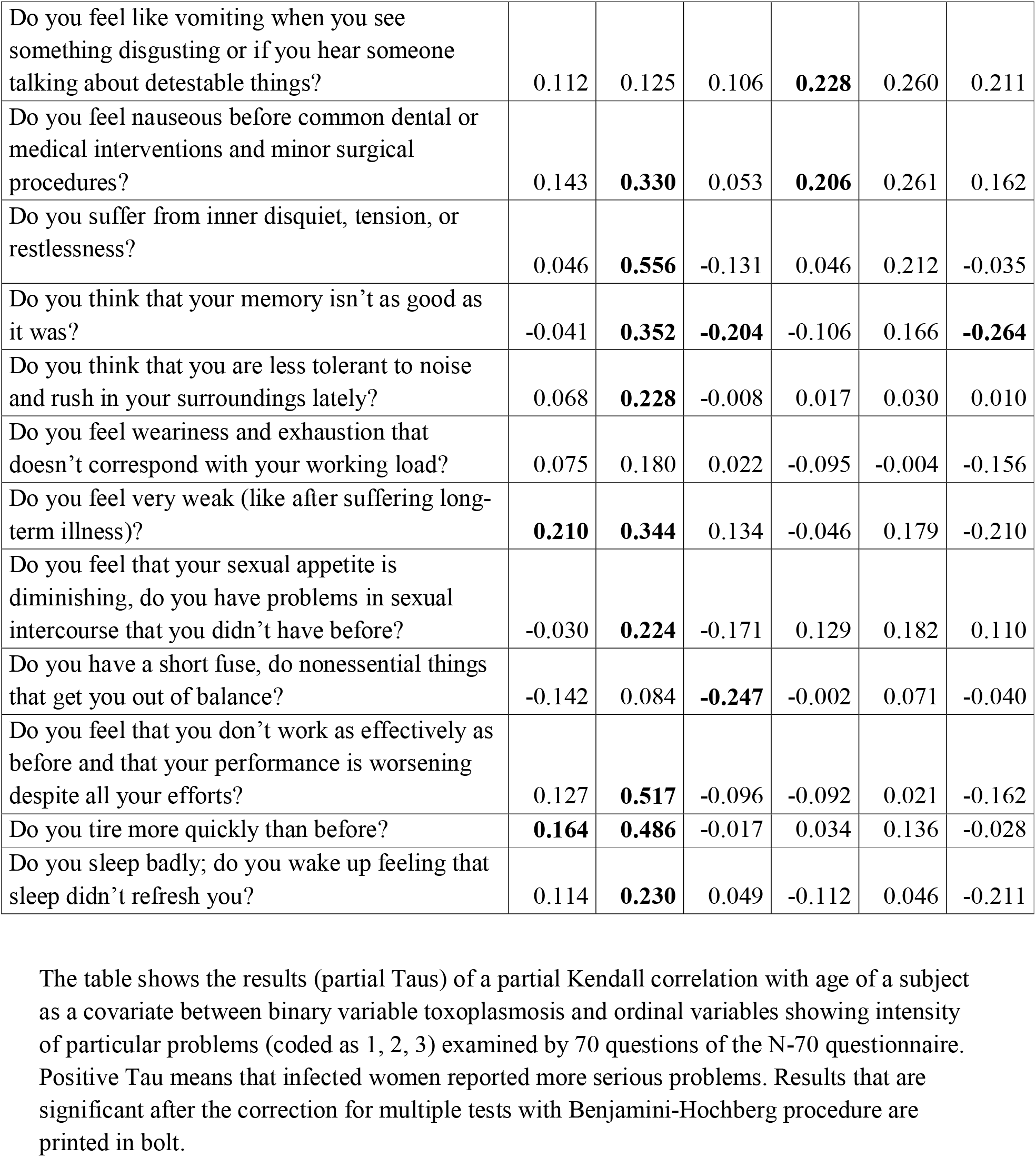
Table 1 Differences between responses of Toxoplasma-infected and *Toxoplasma*-free women to 70 N-70 questions

The existence of a statistical association between two factors does not mean a causal relation between these factors. However, the existence of a causal relation can be supported when one or more Bradford-Hill criteria of causality are fulfilled. To search for such support, we tested the biological gradient of the effect of toxoplasmosis. This was achieved by studying the correlation between the concentration of anti-*Toxoplasma* antibodies and N-70 factors in a subset of *Toxoplasma*-infected women. Fig. 2 and Table 2 show that the CFT titre of antibodies in *Toxoplasma*-infected women correlate with the total N-70 score, as well as with all seven N-70 factors in Rh factor-negative women but not in Rh factor-positive women.

**Fig. 2.**
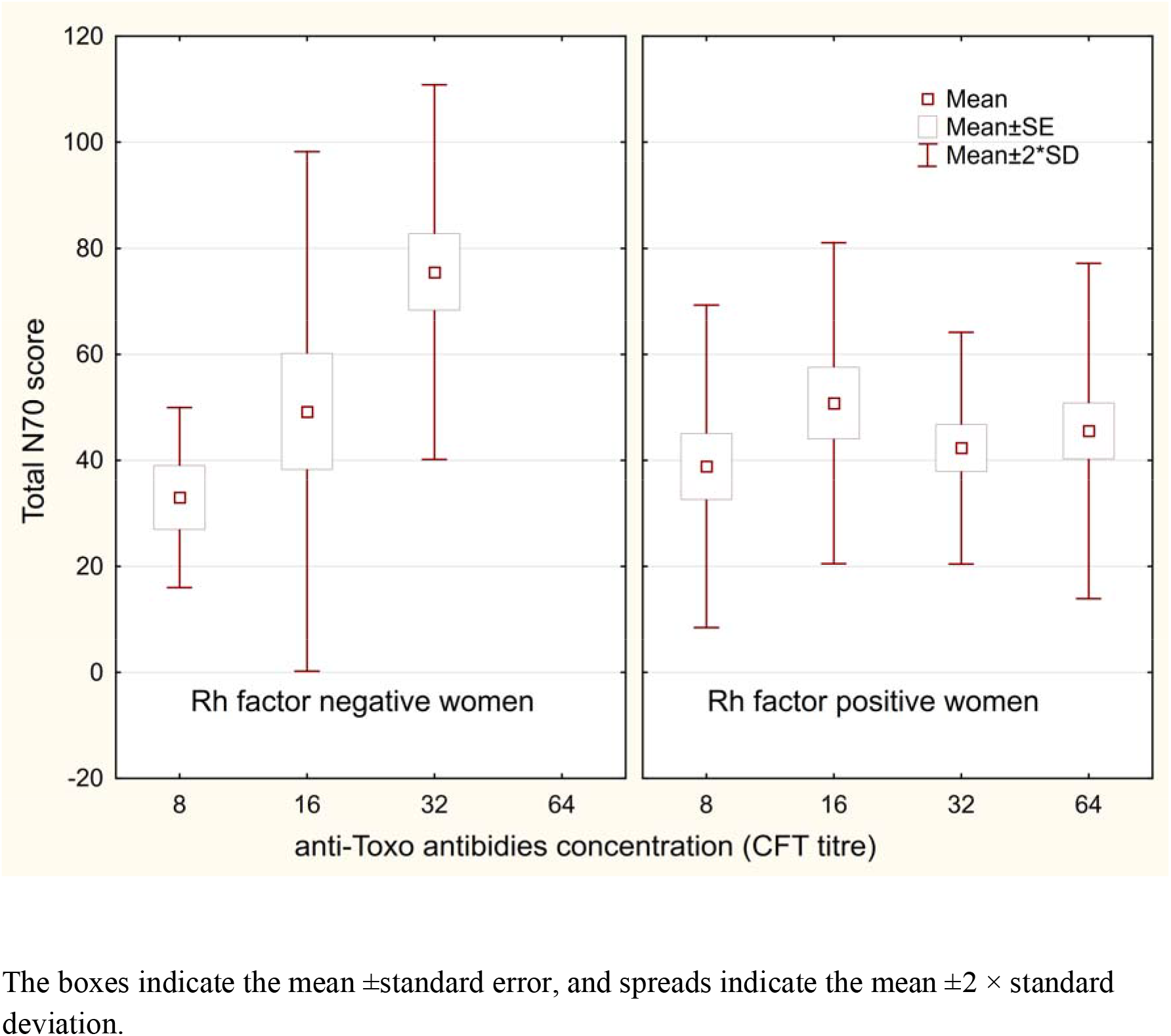
Correlation between N-70 score and anti-*Toxoplasma* antibodies concentration

**Table 2.** Correlation between concentration of anti-*Toxoplasma* antibodies and N-70 factors

|  | New examination |  |  |  | Old examination |  |  |  |
| --- | --- | --- | --- | --- | --- | --- | --- | --- |
|  | Rh negative<br>(N=13) |  | Rh positive<br>(N=26) |  | Rh negative<br>(N=7) |  | Rh positive<br>(N=11) |  |
|  | Tau | p | Tau | p | Tau | p | Tau | p |
| anxita | **0.527** | **0.012** | 0.153 | 0.272 | 0.532 | 0.093 | 0.215 | 0.357 |
| depression | **0.433** | **0.039** | 0.028 | 0.843 | 0.168 | 0.596 | 0.107 | 0.646 |
| obsession | **0.481** | **0.022** | 0.104 | 0.456 | **0.802** | **0.011** | 0.324 | 0.165 |
| histeria | **0.585** | **0.005** | 0.189 | 0.176 | 0.363 | 0.253 | 0.335 | 0.151 |
| hypochondria | **0.517** | **0.014** | 0.154 | 0.271 | 0.432 | 0.173 | 0.253 | 0.280 |
| vegetative lability | **0.508** | **0.016** | -0.076 | 0.587 | 0.276 | 0.384 | 0.171 | 0.465 |
| psychasteny | **0.427** | **0.042** | -0.026 | 0.854 | 0.491 | 0.121 | 0.191 | 0.413 |

Partial Tau correlations (age controlled) that are significant after the correction for multiple tests are printed in bolt.

Twenty (20) Rh-negative women (7 *Toxoplasma*-infected) and 31 Rh-positive women (11 *Toxoplasma*-infected) also completed the questionnaire 3 years earlier, before they knew whether or not they were infected. We repeated all analyses with this smaller data set. No effects of toxoplasmosis or toxoplasmosis-Rh factor interaction were significant after the correction for multiple tests, possibly due to the low number (7) of *Toxoplasma*-infected, Rh negative women, the results not shown. However, a more detailed analysis once again showed that many of the seventy N-70 questions were answered differently by the *Toxoplasma*-infected and *Toxoplasma*-free women (11 in the whole set, 5 in the subset of 20 Rh-negative women and 6 in the subset of 31 Rh positive women; Benjamini-Hochberg correction for multiple tests), see Tab. 1. Fig. 3 shows that N-70 pathognomonic factors mostly decreased or stayed constant between the first and the second examination in Rh-positive *Toxoplasma*-infected and *Toxoplasma*-free women and in Rh-negative *Toxoplasma*-free women, but three factors (anxiety, depression, and psychasthenia) increased in Rh-negative *Toxoplasma*-infected women. The repeat measure ANCOVA with N-70 factors, measured before testing for toxoplasmosis and about 3 years later, showed that the time-toxoplasmosis-Rh interaction had a significant effect only on depression (p=0.027, μ^2^=0.10).

**Fig. 3.**
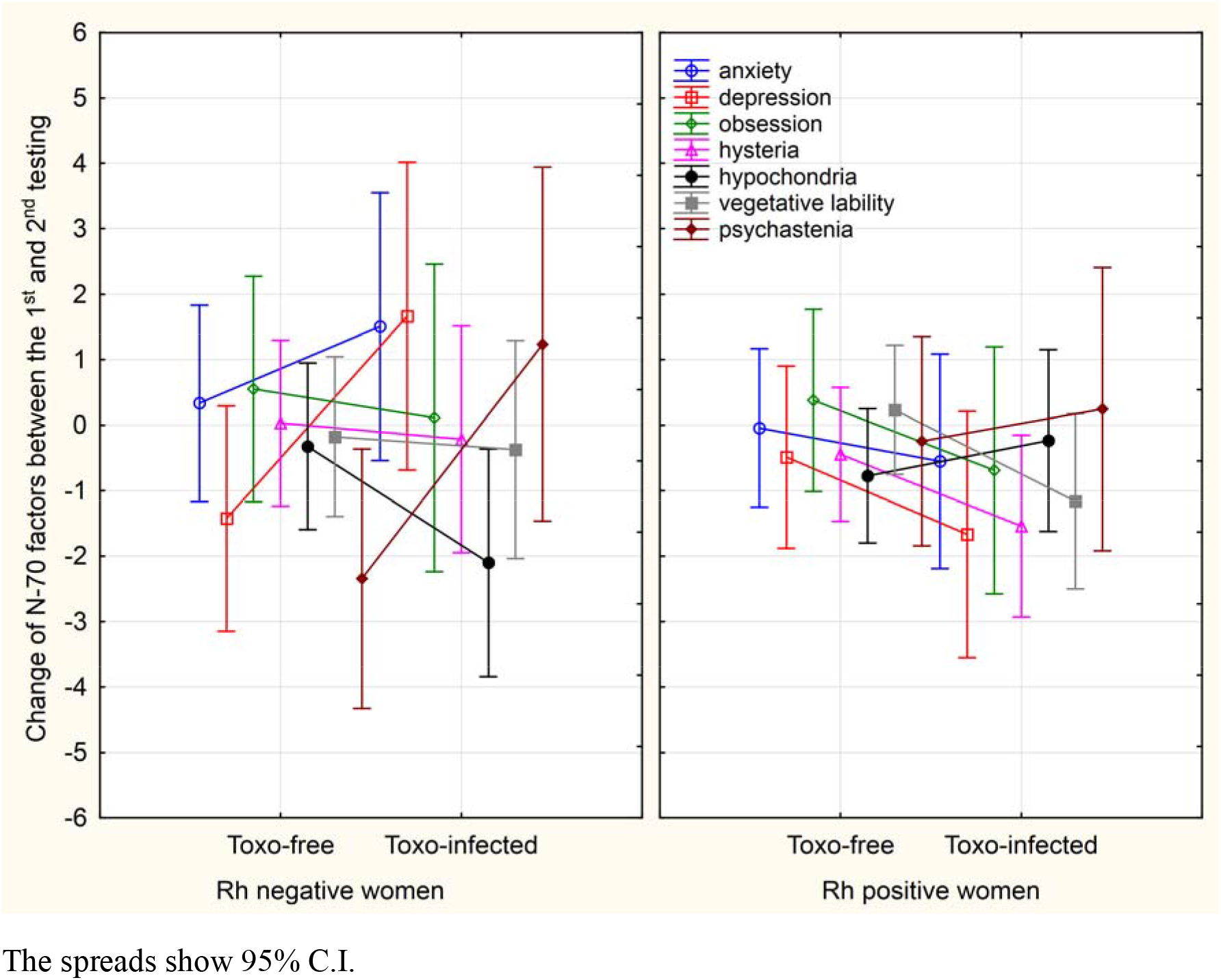
Changes in N-70 factors between the first and second testing

In the anamnestic questionnaire, the subjects rated their health status before testing for toxoplasmosis, and during N-70 testing 3 years later. Tab. 3 shows that infected Rh-negative as well as the Rh-positive women express several indices of impaired physical and mental health before, as well as after, obtaining the information about their toxoplasmosis status. For example, they estimated their probable age of survival to be lower than *Toxoplasma*-free women, reported more frequent hospitalizations in the past 5 years, and to have more serious or more frequent neurological problems.

**Table 3.** Differences in health related variables between *Toxoplasma*-infected and *Toxoplasma*-free women

|  | New examination |  |  | Old examination |  |  |
| --- | --- | --- | --- | --- | --- | --- |
|  | All<br>(N=79) | Rh-<br>(N=27) | Rh+(N=52) | All<br>(N=51) | Rh-<br>(N=20) | Rh+<br>(N=31) |
| allergic problems | 0.05 | **0.23** | -0.06 | 0.14 | 0.21 | 0.12 |
| dermatologic problems | -0.09 | 0.01 | -0.16 | 0.01 | 0.03 | -0.02 |
| digestive organs problems | 0.05 | 0.03 | 0.07 | 0.09 | 0.14 | 0.05 |
| metabolic problems | -0.02 | -0.18 | 0.05 | 0.09 | -0.15 | 0.21 |
| infectious dis. problems | -0.06 | -0.01 | -0.09 | -0.10 | -0.10 | -0.11 |
| cardiovascular problems | 0.07 | 0.00 | 0.09 | -0.17 | 0.00 | -0.21 |
| orthopedic problems | -0.03 | 0.07 | -0.10 | 0.00 | 0.03 | -0.10 |
| neurological problems | 0.13 | **0.23** | 0.09 | **0.25** | 0.26 | **0.26** |
| headache | -0.01 | 0.20 | -0.11 | -0.03 | -0.02 | -0.09 |
| other chronical pain | -0.08 | 0.16 | **-0.18** | 0.15 | 0.07 | 0.20 |
| other chronical problems | 0.04 | 0.06 | 0.03 | 0.15 | -0.05 | **0.26** |
| depression | 0.10 | **0.23** | 0.02 | -0.03 | 0.06 | -0.06 |
| other psychiatric problems | -0.06 | **0.23** | **-0.21** | -0.12 | -0.12 | -0.12 |
| tired after work | 0.06 | **0.29** | -0.06 | -0.02 | 0.14 | -0.18 |
| antibiotics per year | 0.14 | 0.19 | 0.13 | 0.03 | 0.02 | 0.04 |
| doctors' visits per year | 0.08 | 0.04 | 0.10 | 0.16 | -0.06 | **0.37** |
| spending week in hospital | **0.27** | **0.38** | **0.21** | **0.28** | 0.04 | **0.53** |
| life self-expectancy | **-0.24** | **-0.29** | **-0.24** | nd. | nd. | nd. |

Partial Tau correlations (age controlled) that are significant after the correction for multiple tests are printed in bolt. For more details concerning particular health-related variables, see Materials and methods.

## 4. Discussion

The present study showed that *Toxoplasma*-infected women with Rh-negative phenotype expressed higher levels of certain potentially pathognomonic factors measured with the N-70 questionnaire, especially obsessiveness, vegetative lability, and psychasthenia. In the infected subjects, the level of these factors correlated positively with the concentration of anti-*Toxoplasma* antibodies in Rh-negative women but not in more numerous Rh-positive women. Both Rh-negative and Rh-positive women reported more serious and more frequent physical and mental health-related problems. Differences between *Toxoplasma*-infected and *Toxoplasma*-free subjects in potentially pathognomonic factors and health-related variables were also observed in a subset of women who completed the N-70 and anamnestic questionnaires 3 years before the present study, i.e., during a time when they were not aware whether they were infected or not.

Poorer health status of *Toxoplasma*-infected subjects was observed in many case-controlled studies – for a review see (Flegr et al., 2014). A recent study performed on a large nonclinical cohort of 1,486 volunteers showed that *Toxoplasma*-infected subjects scored more poorly in 28 of 29 health-related variables, including number of stays in the hospital, number of doctors visits, frequency of being tired, and seriousness or frequency of allergic, neurological and mental health problems. In contrast to previous studies, here we demonstrated the existence of toxoplasmosis-related effects before the subjects were informed of whether they are *Toxoplasma*-infected or not. This means that the observed effects could not be the result of autosuggestion based on the subjects’ belief that toxoplasmosis has a negative effect on their health. The only factor that significantly increased in Rh-negative, Toxoplasma-infected women was depression, i.e., the factor that was not significantly higher in *Toxoplasma*-infected than in *Toxoplasma*-free Rh-negative women during the second N-70 testing. Our conclusion – that impaired physical and mental health status, not autosuggestion, was responsible for higher N-70 factors – was also supported by the fact that *Toxoplasma*-infected women had the same level of hypochondria as the *Toxoplasma*-free women. The negative effects of toxoplasmosis on human health were also demonstrated in a recent ecological study (Flegr et al., 2014). The prevalence of toxoplasmosis in 88 countries correlated positively with specific disease burden for 23 of 128 studied disorders. The effects of toxoplasmosis on public health was relatively large. For example, the differences in prevalence of toxoplasmosis explained 23 *%* of inter country variability in total disease burden in Europe (Flegr et al., 2014).

At the same time, the present results concerning factors in women measured with the N-70 questionnaire contrasted with results of a similar study performed on male soldiers who went through the entrance examinations for (well-paid) participation in international military missions. The authors of the study suggested that the *Toxoplasma*-infected subjects were objectively motivated to mask their negative and accentuate their positive characteristics. It is, however, also possible that toxoplasmosis had an opposite effect on these pathognomonic factors in men and women as it has been already demonstrated for many personality factors and behavioral patterns (Lindová et al., 2006; Lindová et al., 2010; Flegr et al., 2011).

Both the present and previous studies show a much stronger effect of toxoplasmosis on Rh-negative than Rh-positive subjects. In the present study, we observed significantly higher N-70 factors in *Toxoplasma*-infected Rh-negative women and slightly (not significantly) lower N-70 factors (i.e., better physical and mental health) in *Toxoplasma*-infected Rh-positive women. Detailed analyses of the N-70 questionnaire responses of Toxoplasma-infected and *Toxoplasma*-free Rh-positive women showed that 11 of 12 significant differences indicated better physical or mental health of infected subjects. Such paradoxical (positive) effects of *Toxoplasma* infection have been previously reported to exist. The case control study performed on 500 blood donors showed that *Toxoplasma*-infected Rh-positive heterozygotes showed better psychomotor performance – namely, shorter reaction times – than *Toxoplasma*-free Rh-positive heterozygotes. At the same time, psychomotor performance of infected Rh-positive homozygotes and especially of infected Rh-negative homozygotes was much poorer than that of corresponding controls (Novotná et al., 2008). It has been shown that toxoplasmosis has various effects on human physiology and some of these effects, such as increase of testosterone (Flegr et al., 2008a; Flegr et al., 2008b) or partial immunosuppression (Flegr and Stříž, 2011), could have a positive impact on human performance and health in certain situations. It must be remembered that our population of Rh-positive women represents a mixture of homozygotes and heterozygotes. At the same time, current data suggests that protective effects of Rh factor positivity are fully expressed only in the Rh-positive heterozygotes (Novotná et al., 2008; Flegr, 2016). Therefore, future studies should be performed on DNA genotyped populations.

A combination of results of observational studies performed on humans, and especially experimental studies performed on animals, suggests (but of course does not prove) that toxoplasmosis is the cause rather than the effect of impaired physical and mental health of infected hosts. The present study showed that the level of N-70 pathognomonic factors correlated with the level of anti-*Toxoplasma* antibodies measured with CFT. This test measured the concentration of specific IgM and of certain subclasses of IgG antibodies reacting with various *Toxoplasma* antigens. It has been shown that the intensity of the CFT signal decreases with time from the infection and could be therefore used as a proxy of the duration of latent toxoplasmosis. The intensity of many toxoplasmosis-associated changes, such as personality factor changes (Flegr et al., 2000) or the impairment of reaction times (Havlíček et al., 2001), are negatively correlated with the concentration of anti-*Toxoplasma* antibodies measured with CFT. It has been shown that CFT titres are probably a better proxy for the duration of the infection than the concentration of IgG measured with enzyme-linked immunosorbent assay (Kodym et al., 2007) but worse than – now rarely used – indirect immunofluorescence assay (Kaňková et al., 2007). The existence of a negative correlation is mostly considered to be indirect evidence for models suggesting that given toxoplasmosis-associated changes represent slow cumulative effects of latent infection. In contrast, a positive correlation between the concentration of antibodies and the changes suggests that the observed effects represent transient vanishing effects of acute toxoplasmosis. Such positive correlations were observed, for example, for the probability of traffic accidents (Flegr et al., 2009) and predisposition to gestational diabetes mellitus (Kaňková et al., 2015), and now additionally for N-70 factors.

### Limitations

The main limitation of the present study is that its participants were tested for toxoplasmosis several years before the current study. Toxoplasmosis is a life-long infection and, therefore, women with positive results of the serological test stay infected for their entire lives.

However, at least some originally *Toxoplasma*-free women could have acquired the infection in the years that passed between the diagnosis and the start of the present study. Monte Carlo model-based analysis showed that the presence of such false negative subjects in the study population could easily result in Type 2 error, i.e., failure to detect existing effects of toxoplasmosis, but could not result in false positive results of a study – in the detection of non-existing effects (Flegr and Horáček, 2017).

### Conclusions

Our results support the notion that many behavioral effects of toxoplasmosis represent side-effects of mild but long-term impaired health status rather than a product of adaptive parasitic manipulation aimed to increase the chances for the transmission of *Toxoplasma* from intermediate to definitive hosts. The results also suggest that toxoplasmosis likely has a serious impact on subjectively perceived physical and mental health. Toxoplasmosis affects about one third of the human population worldwide and therefore its global impact on public health may be important. No method of treatment of latent toxoplasmosis and no vaccine that could protect humans against the infection are currently available. The results that slowly accumulated over the past 15 years strongly suggest that such methods and vaccines should be intensively searched for.

## Authors’ contribution

JF designed the study and performed the analyses. BŠ collected and preprocessed the data. Both authors contributed to and have approved the final manuscript.

## Acknowledgement

We would like to thank Charlie Lotterman andLucy Burns for their help with the final version of the paper.

